# Quantitative host-response mapping enables phenotypic profiling of viruses detected by virome sequencing

**DOI:** 10.64898/2026.10.13.758418

**Authors:** Junna Kawasaki, Luca Nishimura, Michiaki Hamada, Edward C. Holmes, Jumpei Ito, Tadaki Suzuki

## Abstract

Animal-derived viruses have repeatedly caused human outbreaks, motivating efforts to identify and characterize viruses circulating in animals. Virome sequencing has accelerated animal virus discovery, but phenotypic characterization remains difficult to scale. Although generated for viral detection, virome RNA-seq data also capture host transcriptional responses to infection. Here, we developed ViroChart, a comparative transcriptomic framework that quantifies activities across biological pathways and maps diverse mammalian host responses into a common, interpretable low-dimensional space. ViroChart enables clustering of infected samples by host response pattern and quantitative assessment of host response similarity between samples. In virome data sets from wild and farmed animals, some lung samples infected with poorly characterized betacoronaviruses showed host responses resembling those induced by viruses capable of causing fatal disease. ViroChart also revealed host-response heterogeneity in natural infections: among rats with zoonotic rodent-borne viruses, most showed weak responses, consistent with asymptomatic infection, but some showed pronounced inflammation characteristic of acute infection. ViroChart thus enables phenotypic characterization and prioritization of newly detected viruses by leveraging the same RNA-seq data from which their genomes were detected.

**Highlights:**

- ViroChart characterizes viruses from host responses in virome RNA-seq data
- Embeds RNA-seq data from diverse hosts and viruses in a shared latent space
- Reveals host-response heterogeneity across naturally infected animals
- Fatal-infection-like responses flag viruses warranting further study

**Graphical Abstact:** 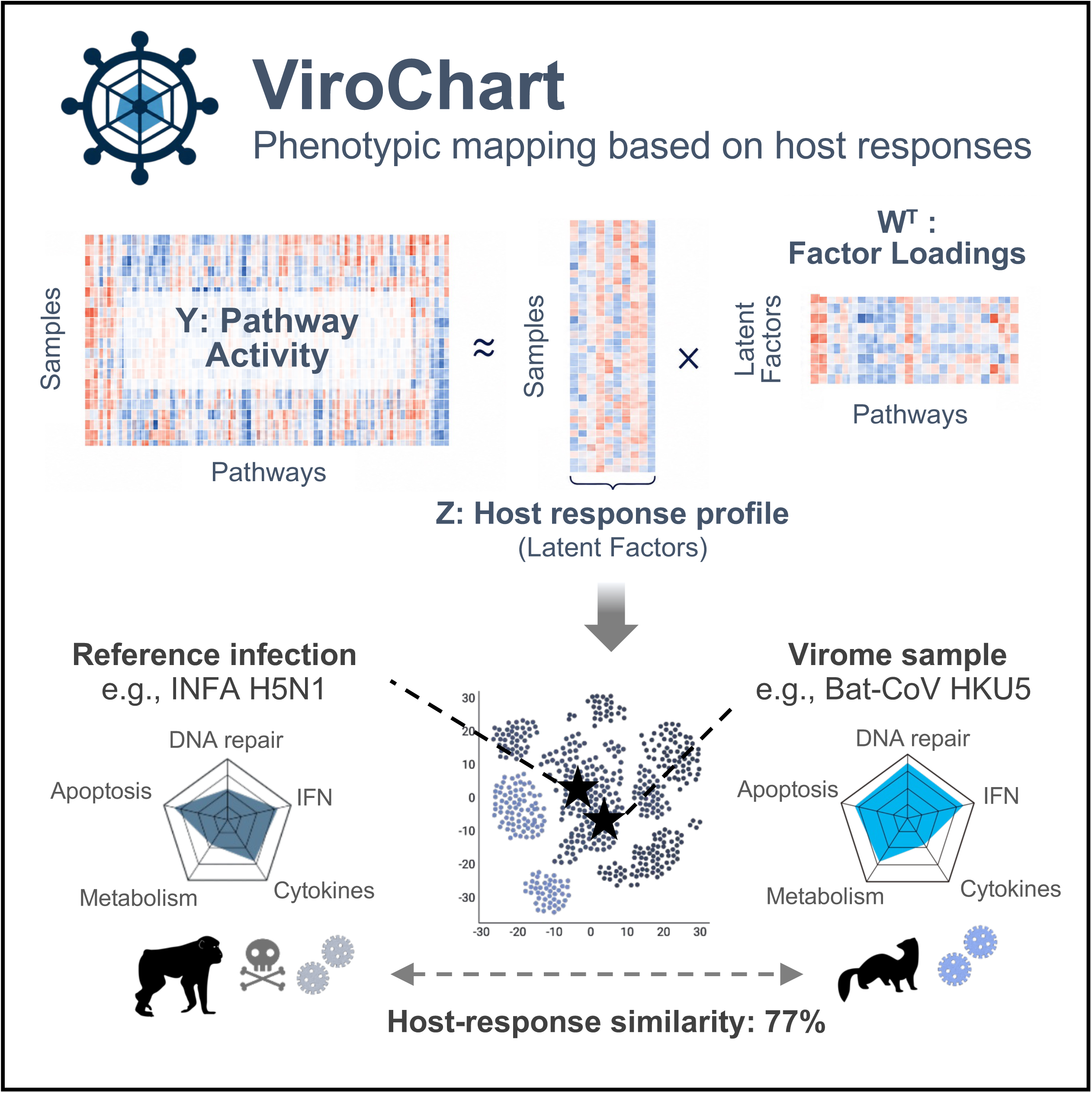

## Introduction

As exemplified by pandemic influenza and COVID-19, animal-derived viruses have repeatedly caused major infectious disease outbreaks in humans^1^. Virome analysis, which enables comprehensive virus detection using high-throughput sequencing, has become a powerful approach for discovering novel viruses in wild and farmed animals that may pose future threats to human health^2–4^. Indeed, approximately half of the novel animal-derived virus species registered by the International Committee on Taxonomy of Viruses (ICTV) since 2000 have been discovered through virome studies^5^. However, the phenotypic characterization of newly identified viruses, including their pathogenicity, still relies heavily on *in vitro* and *in vivo* experiments^3,6^. Such experimental characterization is difficult to scale to the rapidly increasing number of newly discovered viruses. Moreover, suitable experimental systems, such as susceptible cell lines that support viral replication, have not been established for many viruses. Consequently, a growing gap exists between the rapid discovery of viral genomes and the characterization of their biological properties.

This gap may be bridged by the virome data themselves, as RNA-seq–based virome analysis inevitably captures host gene expression profiles alongside viral genomes. Indeed, we previously showed that the expression patterns of interferon-stimulated genes (ISGs), a hallmark of the innate immune response, can serve as informative signals for detecting viral infection^7^. Because host transcriptional signatures change dynamically during infection and reflect biological processes involved in disease pathogenesis^8^, they may be informative not only for detecting infection, but also for inferring the biological characteristics of the infecting virus.

However, comparing host transcriptional responses across virome data sets remains challenging. One issue is data set heterogeneity: host species, sampled tissues, and sampling procedures vary substantially among cohorts, complicating conventional differential gene expression analyses based on predefined group comparisons. A second issue is that viral infection elicits coordinated changes across multiple biological pathways, including innate immunity, inflammatory responses, cell death, and metabolic regulation^9–11^. Capturing these multifaceted responses requires considering the activities of many pathways simultaneously, which yields high-dimensional and potentially noisy profiles that are difficult to compare systematically across samples. We therefore hypothesized that integrating diverse pathway activities into a common low-dimensional representation would allow host responses to be systematically compared across heterogeneous data sets. For newly discovered viruses, comparison of such representations with reference profiles from well-characterized viruses could then provide information about their biological properties.

Herein, we present ViroChart, a comparative transcriptomic framework that integrates host transcriptional responses from diverse animals into a common, interpretable low-dimensional space. ViroChart revealed that some poorly characterized viruses detected in animal virome samples were associated with inflammatory response profiles that resemble those observed during experimental infection with highly pathogenic avian influenza A (H5N1) virus. ViroChart enables quantitative comparison of host responses from heterogeneous RNA-sequencing data sets, providing additional information for the rapid characterization and prioritization of newly discovered viruses.

## Results

### Overview of ViroChart

ViroChart is a comparative transcriptomic framework for quantitatively analyzing host responses to viral infection across heterogeneous RNA-seq data sets and is applicable to 20 mammalian species. ViroChart applies Bayesian factor analysis to activity scores for 484 KEGG pathways, computed by single-sample gene set enrichment analysis (ssGSEA), reducing them to a small number of latent factors that capture coordinated changes across the pathways. Individual RNA-seq samples are thereby embedded in a shared low-dimensional space (**Fig. 1A**).

**Fig. 1.**
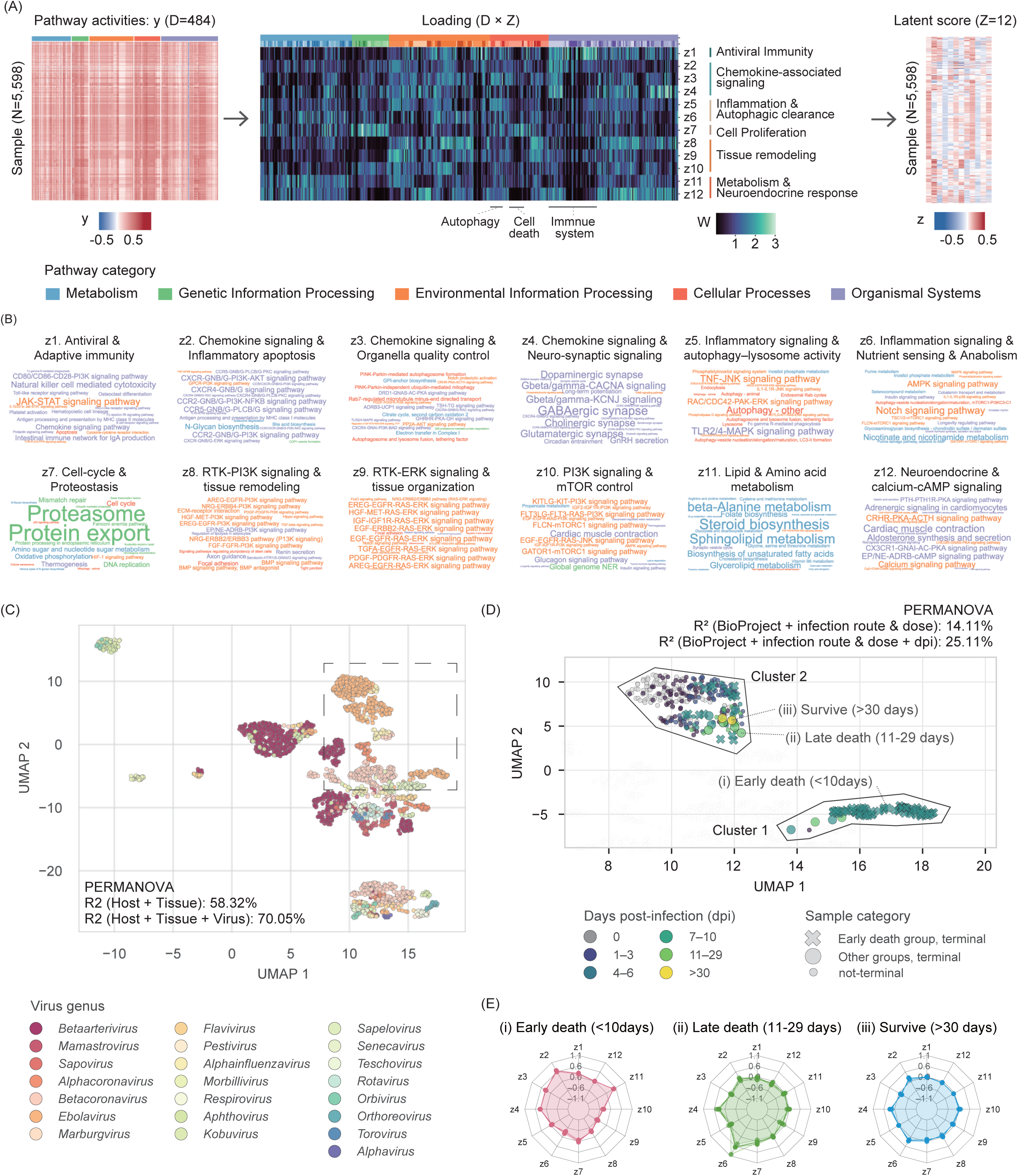
Characterization of host responses by ViroChart. **(A)** Overview of ViroChart. Activity scores (y) for 484 KEGG pathways are mapped to a 12-dimensional vector of latent factor scores (z) using Bayesian factor analysis. The loading matrix W represents the contribution of each pathway to the latent factors. **(B)** Word clouds showing the 20 KEGG pathways with the highest loadings for each latent factor. Font size is proportional to the pathway loading. **(C)** UMAP visualization based on the latent factor scores. Only infected samples are shown, with the exception of pre-infection samples from the Ebola virus infection experiments shown in (D). Points are colored according to the infecting virus genus. The gray dashed box indicates the area enlarged in (D). **(D)** Enlarged UMAP visualization of the Ebola virus infection data sets. Point colors indicate days post-infection (dpi), and point shapes indicate sample category. Pale white points represent samples other than those from the Ebola virus infection data sets shown in (C). Representative samples shown in (E) are highlighted with bold outlines. **(E)** Representative host response profiles from the experimental Ebola virus infection data sets. Latent factor scores were adjusted by subtracting the mean scores of uninfected samples across the three BioProjects.

For the ViroChart training data set, we collected 5,598 publicly available mammalian RNA-seq samples, comprising 2,482 virus-infected and 3,116 uninfected samples (**Fig. S1A and Data set 1**). To capture diverse transcriptional responses associated with viral infection, we selected virus-infected samples identified through previous large-scale analyses of public RNA-seq data sets, primarily focusing on data sets utilizing experimental infections (**details in Methods**)^7,12^. These samples included infections with RNA viruses from 22 genera across 14 families (**Fig. S1B**). For the uninfected samples, we limited the number of samples to a maximum of five per BioProject and 200 per host species to reduce overrepresentation of particular species or research projects. We determined an appropriate dimensionality for the latent space based on the performance of a *k*-nearest neighbors (*k*NN) model for classifying infecting virus genera from the latent factor scores (**Fig. S1C**). Because classification performance largely plateaued at 12 latent factors, we adopted this dimensionality for the final model.

To interpret the 12 latent factors, we examined the 20 KEGG pathways with the highest absolute loadings on each factor in the loading matrix W (**Fig. 1B and Data set 2**). Latent factors z1–z6 were primarily associated with innate immune and inflammatory processes, including antiviral immunity, chemokine-associated signaling, and inflammatory cytokine signaling. In contrast, z7 was associated with cell proliferation, z8–z10 with tissue remodeling, z11 with metabolism, and z12 with neuroendocrine stress responses. Thus, ViroChart summarized diverse host responses beyond canonical antiviral responses into a set of biologically interpretable latent factors. We therefore used the 12-dimensional vector of latent factor scores as a quantitative representation of each sample’s host response, hereafter referred to as the host response profile.

### ViroChart enables the projection of new data sets into a shared latent space

The trained ViroChart model has a function to project newly acquired RNA-seq data into the existing latent space. To evaluate this capability, we constructed five hold-out models, each excluding a subset of BioProjects (**Fig. S2A**). Samples excluded from each training data set were then projected into the corresponding latent space. The hold-out models showed goodness-of-fit, as measured by log likelihood, comparable to that observed for the full model (**Fig. S2B**). Global and local structures were largely preserved between the hold-out and full-model spaces, as indicated by highly correlated pairwise Euclidean distances among the hold-out samples (**Fig. S2C**) and the substantial agreement of their *k*NN sets (median Jaccard index > 0.60 for all evaluated values of k; **Fig. S2D**). Furthermore, *k*NN classification of infection status and infecting virus genus performed comparably when based on latent factor scores from the full and hold-out models (**Fig. S2E**). Together, these results indicate that ViroChart can accurately project unseen RNA-seq samples into the latent space.

We next assessed the sensitivity of latent factor interpretation to differences in training data set composition. Because independently trained models do not necessarily share the same ordering of latent factors, we constructed 12 × 12 correlation matrices between their loading vectors of the full and hold-out models and applied the Hungarian algorithm to identify the optimal one-to-one matching of factors (**Fig. S2F**). The mean Spearman correlation between matched loading vectors was 0.66 (SD = 0.11) across the hold-out models (**Fig. S2G**). Thus, although some latent factors were affected by training data set composition, pathway loading patterns were largely preserved for many factors, supporting the robustness of their pathway-level interpretation.

### ViroChart captures host-response variation across viral genera and infection states

To examine the structure of the latent space learned by ViroChart in detail, we visualized the latent factor scores using UMAP (**Fig. 1C**). Infected samples broadly clustered according to the genus of the infecting virus. Because host responses are expected to vary across host taxa and tissues, we evaluated the contribution of infecting virus genus after accounting for these factors. Permutational Multivariate Analysis of Variance (PERMANOVA) showed that host genus and tissue together explained 58.32% of the variance in the latent space, while the infecting virus genus explained an additional 11.73% (P = 0.01). Thus, the infecting virus genus explained approximately 28.14% (11.73/41.68%) of the variance remaining after accounting for host genus and tissue. Furthermore, in *k*NN classification of infecting virus genera, classifiers based on the latent factor scores performed comparably to those based on the original 484-dimensional pathway activity scores (median macro-F1 score: latent factor scores, 0.32; pathway activity scores, 0.38; **Fig. S2E**). These results suggest that ViroChart captures differences in host responses to infection across viral genera.

To examine whether ViroChart can capture the temporal dynamics of host responses during infection, we next analyzed longitudinal whole-blood RNA-seq data from three independent experimental Ebola virus infection studies in macaques (*Macaca mulatta* or *Macaca fascicularis*) (PRJNA476001, PRJNA407110, and PRJNA388501; **Data set 3**)^13–15^. These data sets included samples collected up to 41 days post-infection (dpi). Based on survival outcomes, animals were classified into the Early death group (died by 10 dpi), Late death group (died at 11–29 dpi), or Survive group (survived through 41 dpi). In the UMAP, samples were distributed largely according to dpi (**Fig. 1D**). PERMANOVA showed that variation associated with dpi accounted for 11.0% of the variance in the latent space after accounting for BioProject and infection conditions, including the route and dose of virus inoculation (P = 0.01). Thus, ViroChart captures changes in host state over the course of infection.

Furthermore, the Ebola virus-infected samples formed two major clusters in the UMAP: Cluster 1 was enriched for terminal samples (the last samples collected before death or euthanasia at a humane endpoint) from animals in the Early death group, whereas most other samples into Cluster 2 (**Fig. 1D**). Terminal samples from the Early death group showed particularly high z2–z4 scores associated with chemokine responses (**Fig. 1E**), consistent with the enhanced chemokine and inflammatory cytokine responses previously reported in lethal Ebola virus infection^16^. In contrast, terminal samples from the Late death group and end-of-study samples from Survive groups did not show elevated z2 or z3 scores, suggesting host response profiles distinct from those of the Early death group. Together, these results show that ViroChart captures host response features associated with infection progression and disease outcome.

### Systematic characterization of host-response diversity across respiratory virus infections

We next systematically characterized the diversity of host response profiles associated with *in vivo* infections with well-characterized respiratory viruses. We analyzed 317 lung RNA-seq samples comprising infections with four respiratory viruses—SARS-CoV-2, highly pathogenic avian influenza A (H5N1) virus (INFA H5N1), porcine reproductive and respiratory syndrome virus, and bluetongue virus—across five host species (**Data set 4**). Because host species and tissue substantially influenced the ViroChart latent space (**Fig. 1C**), we adjusted the host response profile of each sample by subtracting the mean latent factor scores of uninfected samples from the same host species and tissue. Hereafter, we refer to these adjusted scores as baseline-adjusted host-response profiles. The 317 infected samples were subsequently grouped into 16 clusters by applying *k*-means clustering to the baseline-adjusted host-response profiles (**Fig. 2A–B**). Eleven of these clusters contained samples from multiple BioProjects, suggesting that similar host responses were observed across independent research projects. Furthermore, host responses from the same virus–host combination were sometimes assigned to different clusters, indicating heterogeneity in host responses within the same combination.

**Fig. 2.**
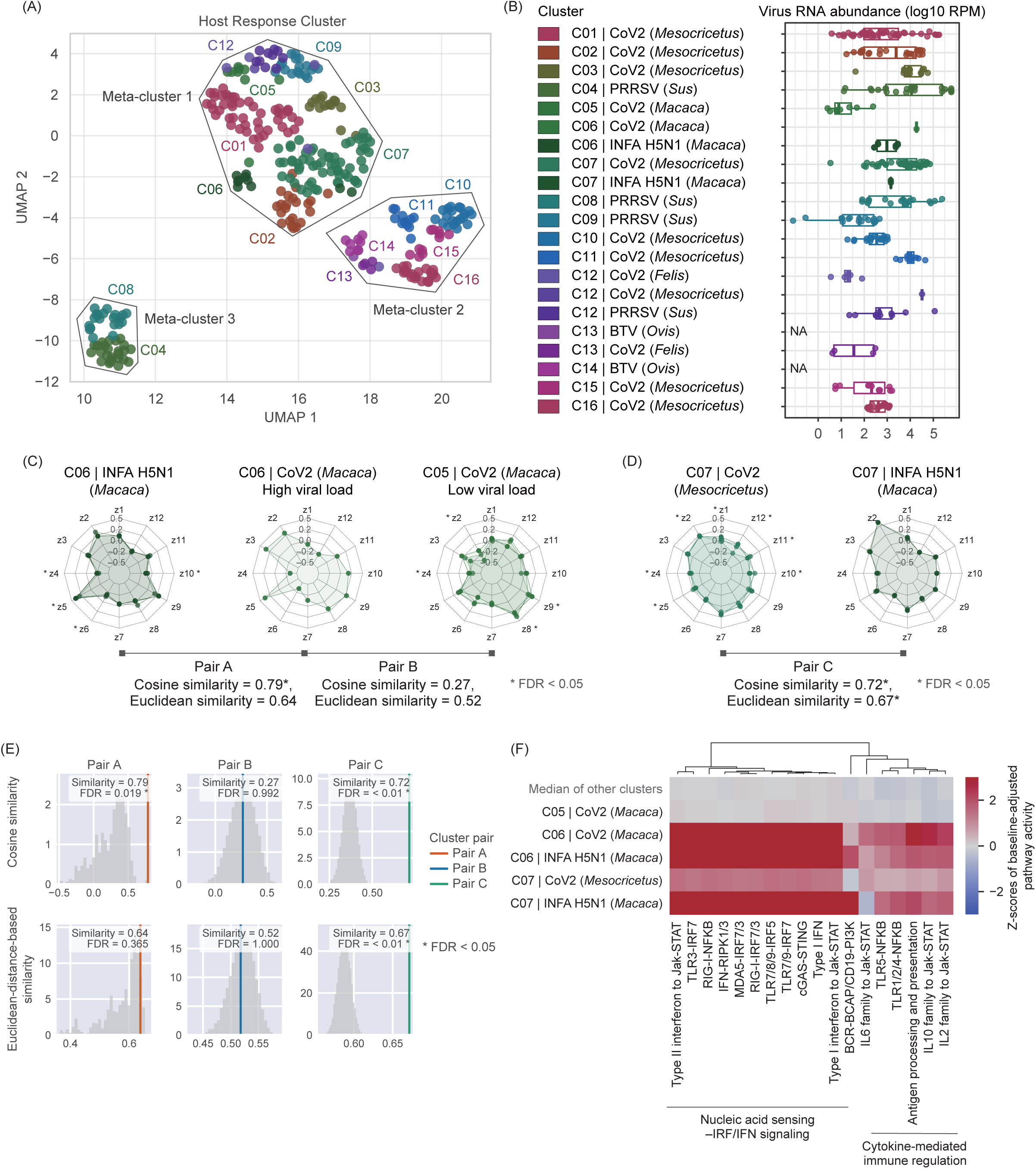
Diversity and similarity of host-response profiles across infections with known respiratory viruses. **(A)** UMAP visualization of respiratory virus-infected samples. Points are colored by unique combinations of host response cluster, virus taxonomy, and host genus. Virus abbreviations are as follows: CoV2, SARS-CoV-2; INFA H5N1, highly pathogenic avian influenza A(H5N1) virus; PRRSV, porcine reproductive and respiratory syndrome virus; and BTV, bluetongue virus. **(B)** Distribution of viral RNA abundance across these combinations. **(C–D)** Examples of similar host responses observed across different host genera or infecting viruses. Radar charts show baseline-adjusted host-response profiles for representative samples. Asterisks indicate latent factors that differed significantly from uninfected samples of the same host genus (Welch’s t-test, Benjamini–Hochberg FDR < 0.05). Cosine similarity and Euclidean-distance-based similarity, as evaluated in (E), are also shown. **(E)** Host-response similarities for the pairs of clusters, evaluated using cosine similarity and Euclidean-distance-based similarity. Histograms show the query-fixed null distributions, and vertical lines indicate the observed mean similarities between clusters shown in (C) and (D). FDR values were calculated from empirical P values using the Benjamini–Hochberg procedure. **(F)** Heatmap of pathway activities related to innate immune and inflammatory responses. These pathways were selected from the KEGG “Immune system” category based on significantly higher activity in C06 or C07 than in the remaining clusters (Benjamini–Hochberg FDR < 0.05). Activities are shown as median z-scores of baseline-adjusted pathway activity for each cluster.

We next examined host response variation within a single virus–host combination, focusing on SARS-CoV-2-infected hamster (*Mesocricetus auratus*) samples, which represented the most abundant combination in our data set. In Meta-cluster 1 (including clusters C01, C02, C03, and C07), latent factor z8, associated with regenerative tissue remodeling, was elevated during the early phase of infection (2–4 dpi), followed by an increase in z2, associated with chemokine-driven immune cell recruitment (**Fig. S3A–C**). These patterns were consistent with temporal changes in host responses reported over the course of SARS-CoV-2 infection^17,18^. In contrast, Meta-cluster 2 (including C10, C11, C15, and C16) showed lower levels of the tissue remodeling-associated z8 and the inflammation-associated factors z5 and z6, representing a distinct host response pattern (**Fig. S3D**). Notably, Meta-cluster 2 contained a high proportion of samples from animals that had received vaccination or therapeutic drug treatment, consistent with the weaker inflammatory responses observed in this group (**Fig. S3A and S3E**)^19–22^. Together, these results show that ViroChart enables systematic comparison of diverse host responses across multiple studies and captures variation associated with biological and experimental characteristics of infection.

### ViroChart identifies shared inflammatory responses across taxonomically distinct viruses

Interestingly, similar host response profiles were observed across infections with taxonomically distinct viruses. For example, cluster C06 contained samples from both INFA H5N1 and SARS-CoV-2 infections in macaques. Most SARS-CoV-2-infected macaque samples were assigned to cluster C05, whereas a subset with high viral RNA abundance was assigned to C06 and showed host response profiles similar to those observed in H5N1 infection (**Fig. 2B–C**).

To quantify these similarities, we used two complementary metrics in the 12-dimensional latent space: cosine similarity, which compares the pattern of the response irrespective of its magnitude, and Euclidean-distance-based similarity, which reflects similarity in both pattern and magnitude. INFA H5N1- and SARS-CoV-2-infected macaque samples in C06 showed high similarity by both metrics, although only the cosine similarity was significant after multiple-testing correction (Pair A in **Fig. 2E**). In contrast, SARS-CoV-2-infected macaque samples in different clusters (C05 and C06) were not significantly similar to each other by either metric (Pair B in **Fig. 2E**). SARS-CoV-2-infected macaque samples in C05 showed relatively high z8 and z9 scores, associated with tissue remodeling, whereas those in C06 showed marked increases in z2 and z5, associated with chemokine-driven immune cell recruitment and inflammatory cytokine responses, respectively **(Fig. 2C)**. Examination of the pathway activities used as input to ViroChart further revealed broad activation of pathways related to innate immune and inflammatory responses in C06 (**Fig. 2F**). Consistent with these patterns, the original INFA H5N1 infection study reported strong inflammatory cytokine responses followed by pyroptosis^23^.

Similar host response profiles were also observed across infections that differed in both the infecting virus and host species **(Fig. 2D)**. INFA H5N1-infected macaque and SARS-CoV-2-infected hamster samples assigned to C07 were significantly similar by both metrics (Pair C in **Fig. 2E**). These samples showed high z2 scores associated with chemokine-driven immune cell recruitment and broad activation of pathways related to innate immune and inflammatory responses **(Fig. 2F)**. Together, these findings show that ViroChart can identify shared inflammatory response patterns across taxonomically distinct viruses, even when infections occur in different host species.

### ViroChart flags poorly characterized viruses with host responses resembling those induced by known pathogenic viruses

We next analyzed host response profiles in 199 lung RNA-seq samples from animal virome studies, which aimed to identify viruses naturally infecting wild and farmed animals through untargeted metagenomic sequencing (**Data set 5**). These samples were obtained from two virome studies (PRJCA026706 and PRJNA1238520) encompassing mammalian species from Carnivora, Rodentia, and Lagomorpha^24,25^. Because these samples were not included in the ViroChart training data set, we projected them into the pretrained latent space and calculated baseline-adjusted latent factor scores. We then mapped the resulting profiles onto the UMAP space constructed from the reference panel of samples infected with known respiratory viruses in **Fig. 2** (**Fig. 3A**).

**Fig. 3.**
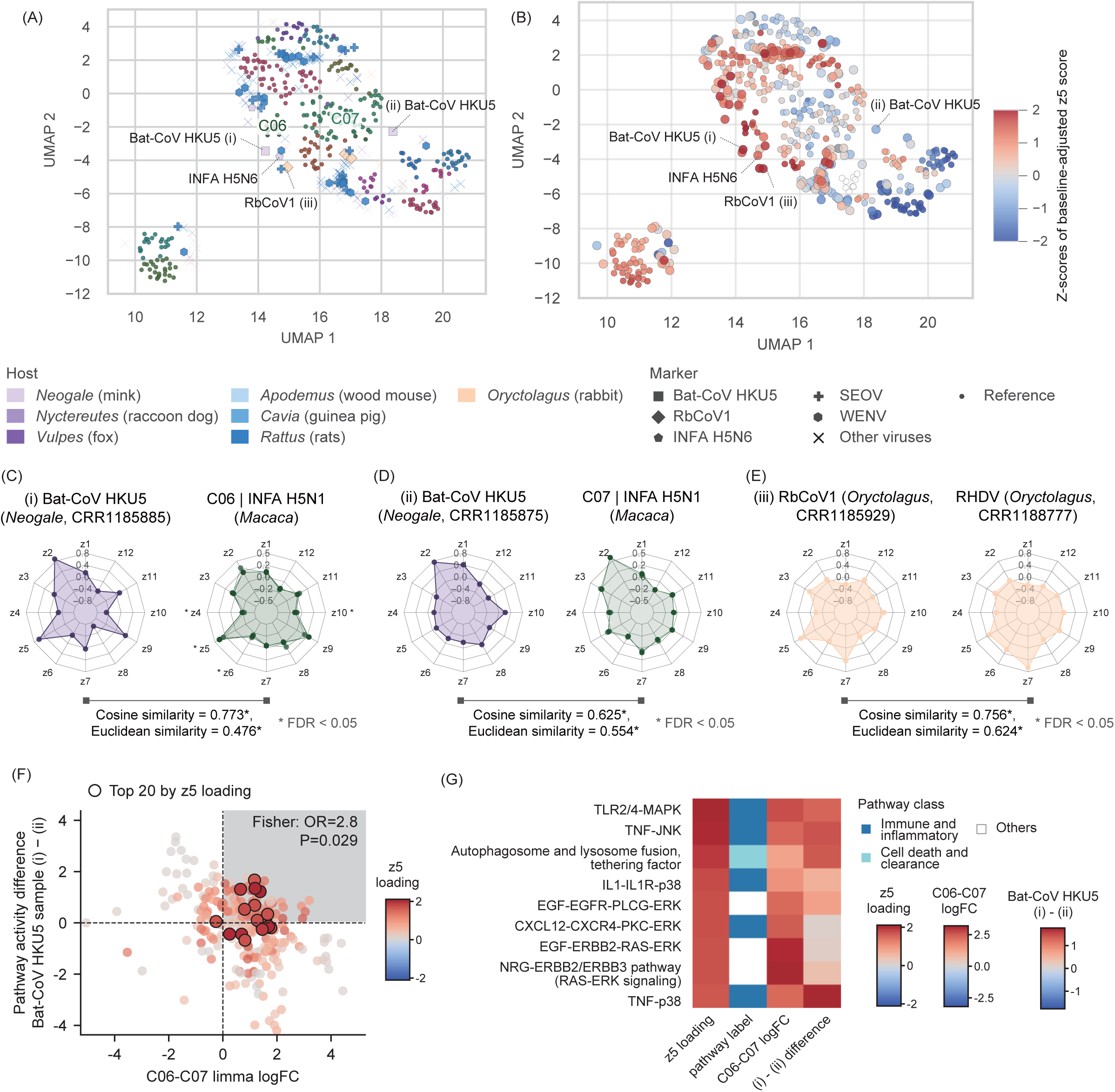
Host response profiles in animal virome data sets. **(A–B)** UMAP visualization of virome samples projected onto the space constructed from the well-characterized respiratory virus infections in Fig. 2A. (i)–(iii) indicate the representative samples shown in (C)–(E). In (A), reference samples are colored by host response clusters, and virome samples are colored by host species. Point shapes indicate the virus species detected by virome analysis. In (B), points are colored by z-scores of baseline-adjusted z5 score. Small points represent reference samples and large points represent virome samples. **(C–E)** Host response profile comparisons between selected virome samples and reference virus-infected samples: Bat-CoV HKU5 compared with C06 (C), Bat-CoV HKU5 compared with C07 (D), and RbCoV1 compared with RHDV (E). Similarity between the indicated sample pairs was evaluated using cosine similarity and Euclidean-distance-based similarity. Statistical significance was assessed by query-fixed permutation testing, with FDR correction using the Benjamini–Hochberg procedure. **(F)** Comparison of pathway activity differences between clusters C06 and C07 and between the Bat-CoV HKU5-positive samples. Points represent pathways and are colored by their z5 loadings. Black outlines indicate the 20 pathways with the highest z5 loadings. Enrichment of these pathways in the upper-right quadrant was evaluated using Fisher’s exact test. **(G)** Pathways in the upper-right quadrant of (F) that were among the 20 pathways with the highest z5 loadings.

As described in **Fig. 2C**, cluster C06 is characterized by strong inflammatory responses and contains samples from INFA H5N1 and SARS-CoV-2 infections. We therefore examined whether any of the virome samples showed host response profiles resembling those of samples in C06. Notably, several virome samples mapped near C06 and showed high z5 scores associated with inflammatory responses (**Fig. 3B**). These samples included minks (*Neogale vison*) infected with highly pathogenic avian influenza H5N6 virus or Pipistrellus bat coronavirus HKU5 (Bat-CoV HKU5; **Fig 3C–D**), as well as rabbits (*Oryctolagus cuniculus*) infected with Rabbit coronavirus 1 (RbCoV1; **Fig 3E**). Examination of the pathway activity scores further showed marked activation of pathways related to innate immune and inflammatory responses in these infections, although the overall response patterns differed among viruses (**Fig. S4A**).

Bat-CoV HKU5 has attracted attention as a virus of potential concern because it caused a pneumonia outbreak in farmed mink following cross-species transmission from bats^24^. Analysis of available data from two Bat-CoV HKU5-positive mink lung samples revealed that these samples exhibited distinct host response profiles. As described above, one infected sample exhibited a high z5 score and showed significant similarity to INFA H5N1-infected samples assigned to C06 (**Fig. 3C**). The other sample exhibited a high z2 score associated with chemokine-driven immune cell recruitment and showed significant similarity to INFA H5N1-infected samples assigned to C07 (**Fig. 3D**). These results suggest that Bat-CoV HKU5-infected samples display host response heterogeneity paralleling that observed among INFA H5N1-infected samples.

To clarify how the transcriptome signatures of the two HKU5-infected mink samples differed, we examined pathways with differential activity between C06-like and C07-like samples. As expected from the markedly higher z5 score in C06, the overlapping pathways—those specifically elevated in reference-panel samples assigned to C06, and also elevated in the C06-like HKU5-infected sample—were enriched for pathways with high z5 loadings (OR = 2.80, P = 0.03) (**Fig. 3F**). These z5-associated pathways mainly governed immune and inflammatory signaling and cell death and clearance, suggesting that the HKU5-infected mink in C06 mounted stronger inflammatory and cytotoxic responses than the mink in C07 (**Fig. 3G**).

RbCoV1 provided another example of host-response profiling for a poorly characterized virus. This virus was identified as a novel betacoronavirus in the original study, but little pathological or epidemiological information is available^24^. RbCoV1-infected samples were assigned to C06, with host response profiles highly similar to those observed during Rabbit hemorrhagic disease virus (RHDV) infection, which causes lethal disease in rabbits^26^ (**Fig. 3E**). Together, these examples show that ViroChart can characterize poorly understood viruses by comparing their host responses with those of well-characterized virus infections. Such comparisons may help prioritize newly detected viruses for further investigation, particularly when little is known about their biological properties (see **Discussion**).

### Natural infection samples do not always exhibit strong host responses

We next quantified the magnitude of host responses in naturally infected samples from virome data sets. For each virus-infected sample, we quantified host response intensity as the sum of the absolute baseline-adjusted latent factor scores. Response intensity was then converted to an uninfected-reference z-score, with the reference distribution defined by uninfected samples from the same host genus (**Fig. 4**). For this analysis, we included RNA-seq samples from multiple organs, covering infections with viruses from 25 families. To increase host diversity, we included two additional BioProjects from bats (PRJNA1184956 and PRJNA929070)^27,28^, resulting in a total of 1,002 RNA-seq samples (**Data set 6**). Among virus-positive samples, 27.25% had uninfected-reference z-score of response intensity greater than 1, compared with 24.14% of the uninfected reference samples (Fisher’s exact test; odds ratio = 1.18, P = 0.31). These results suggest that samples exhibiting host gene expression patterns distinct from those of uninfected samples were relatively uncommon.

**Fig 4.**
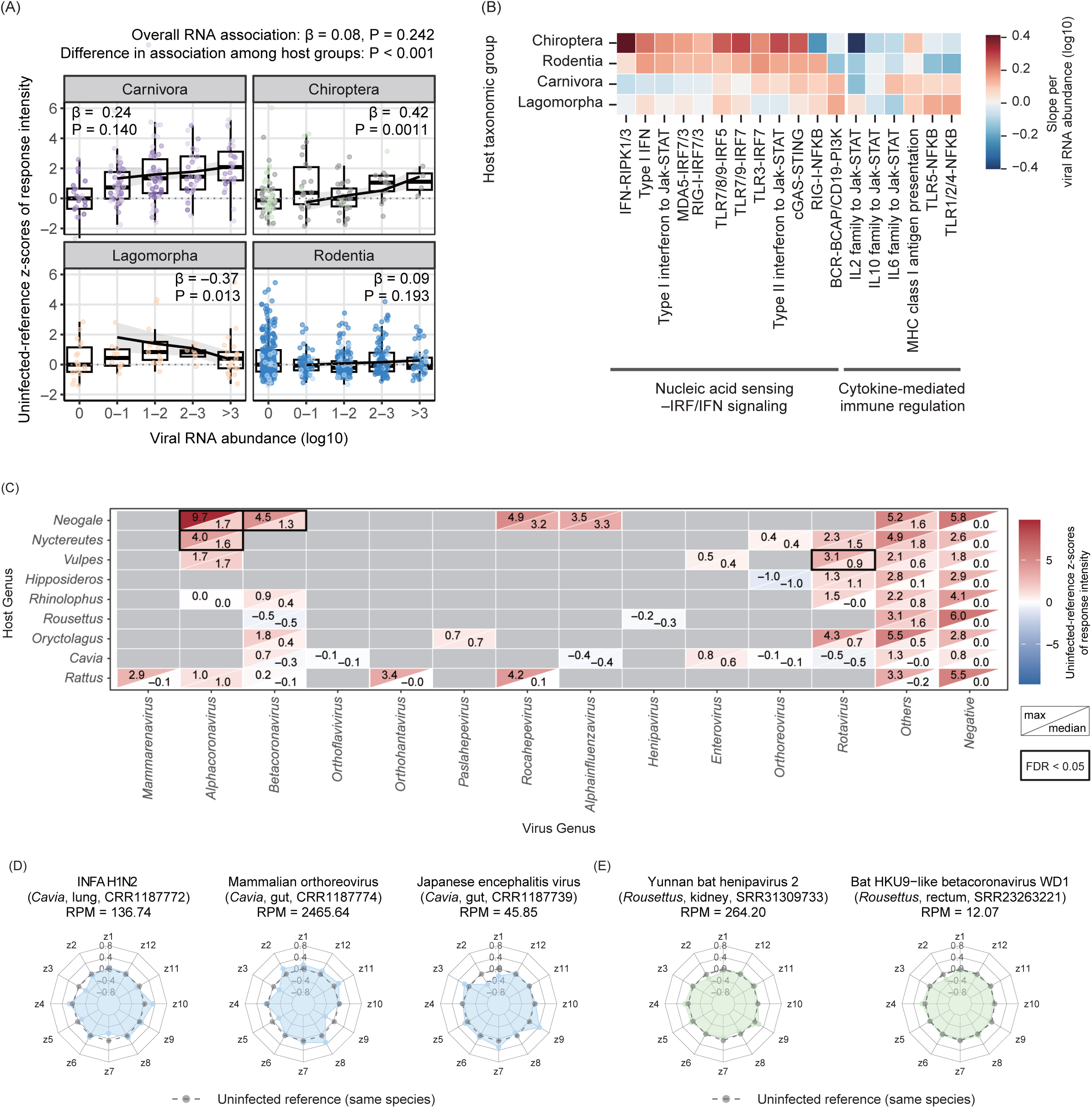
Host response intensity in virome data sets. **(A)** Association between viral RNA abundance and host response intensity. Viral RNA abundance is shown as log10(RPM + 1). All regression analyses included only samples with viral RNA abundance >10 RPM. The overall association was estimated using a model adjusted for host taxonomic group and infecting virus family. Host-group-specific associations were estimated using the interaction model, with the overall interaction evaluated using a Wald test. Regression lines show the estimated association for each host taxonomic group, with shaded areas representing 95% confidence intervals. (B) Host-group-specific associations between viral RNA abundance and the activities of pathways related to innate immunity and inflammation. The pathways shown are the same as those in Fig. 2F. The pathway-specific uninfected-reference z-scores were analyzed using the interaction model described in (A). Colors indicate the estimated slopes for each host taxonomic group and pathway. No association remained significant after Benjamini–Hochberg correction. **(C)** Heatmap of host response intensity across virus–host combinations. For each combination, the upper-left and lower-right triangles show the maximum and median host response intensity, respectively. Bold outlines indicate virus–host combinations with adjusted response intensities significantly greater than zero (one-sided tests with Benjamini–Hochberg correction; FDR < 0.05). **(D–E)** Representative virome samples with particularly weak host responses. Radar charts show their baseline-adjusted host-response profiles. The gray dashed line in each chart shows the mean profile of uninfected samples from the same host species.

To investigate the biological factors underlying variation in the magnitude of host responses, we examined its association with viral RNA abundance. Host response intensity tended to increase with viral RNA abundance after accounting for infecting virus families and host taxonomic groups (**Fig. 4A**). However, the strength of this association differed significantly among host taxonomic groups (Wald test for interaction, P < 0.05). A clear positive relationship was only observed in chiropterans, whereas the relationship was weaker in the other host groups.

We next assessed the association between viral RNA abundance and pathway-level host response intensity. Although none of these associations remained significant after multiple-testing correction, their association patterns differed among host taxonomic groups (**Fig. 4B and Fig. S4B**). In Chiroptera and Rodentia, relatively strong positive associations were observed primarily for pathways related to nucleic acid sensing–IRF/IFN signaling. Conversely, pathways related to cytokine-mediated immune regulation showed no comparable positive trend in these animal groups. These trends were broadly similar to previous observations in some bat and rodent reservoir hosts, where antiviral responses can be induced while excessive inflammatory responses are limited^29–31^.

We next examined differences in host response intensity across virus–host combinations (**Fig. 4C**). Although host response intensity tended to be positive across most combinations, the magnitude of these responses was generally modest. After adjusting for viral RNA abundance, only a limited number of virus–host combinations showed response intensities significantly above the uninfected reference (one-sided tests with Benjamini–Hochberg correction; FDR < 0.05). Notably, several novel virus–host associations involving *Cavia* were reported in the original study^24^, including H1N2 influenza A virus (*Alphainfluenzavirus*), Japanese encephalitis virus (*Orthoflavivirus*), and mammalian orthoreovirus (*Orthoreovirus*), and these infected samples showed weak host responses despite relatively high viral RNA abundance (**Fig. 4D**). Similar weak responses were observed in *Rousettus* samples positive for viruses belonging to *Henipavirus* or *Betacoronavirus* (**Fig. 4E**). Hence, these examples show that high viral RNA abundance does not necessarily coincide with a strong host response in naturally infected animals, although the underlying mechanisms remain unclear (see **Discussion**).

### ViroChart reveals heterogeneous host responses to rodent-borne viruses, including strong inflammation

We next examined host-response heterogeneity within each of two rodent-borne virus infections. Seoul orthohantavirus (SEOV) and Wenzhou arenavirus (WENV) are zoonotic viruses of public health concern that are generally considered to establish asymptomatic infections in their natural rodent hosts^32–34^. The virome data set included 37 lung samples from rodents infected with SEOV or WENV, providing an opportunity to examine host-response variation during natural infection (**Data set 7**). In the UMAP projection, SEOV- and WENV-infected samples were widely distributed, indicating substantial variation in host response profiles even among samples infected with the same virus species (**Fig. 3A**). PERMANOVA showed no significant association between host response profiles and the variables examined, including virus species, viral RNA abundance, the number of coinfecting virus families, and host species.

Previous experimental studies of SEOV have shown that infection of rat cells induces strong IFN responses during early infection (e.g., 4 dpi), followed by a shift toward a more immunotolerant state at later stages (e.g., 12 dpi)^35^. To examine whether the host response diversity observed in naturally infected animals resembled this temporal variation, we analyzed the virome data set described above together with an SEOV *in vitro* infection data set sampled at 4 and 12 dpi (PRJNA1030382; **Fig. 5A–B**). Approximately half of the naturally infected samples showed little activation of innate immune and inflammatory pathways, consistent with largely asymptomatic infection in natural hosts. In contrast, a subset showed responses comparable to or stronger than those observed at 4 dpi in experimental SEOV infection. Notably, these strong responses were observed not only in SEOV-but also in WENV-infected samples, and some samples showed host response profiles significantly similar to those of INFA H5N1-infected macaques assigned to C06 (**Fig. 5C**). Together, these results show that ViroChart reveals host-response heterogeneity among naturally infected animals, from weak responses to pronounced inflammation.

**Fig 5.**
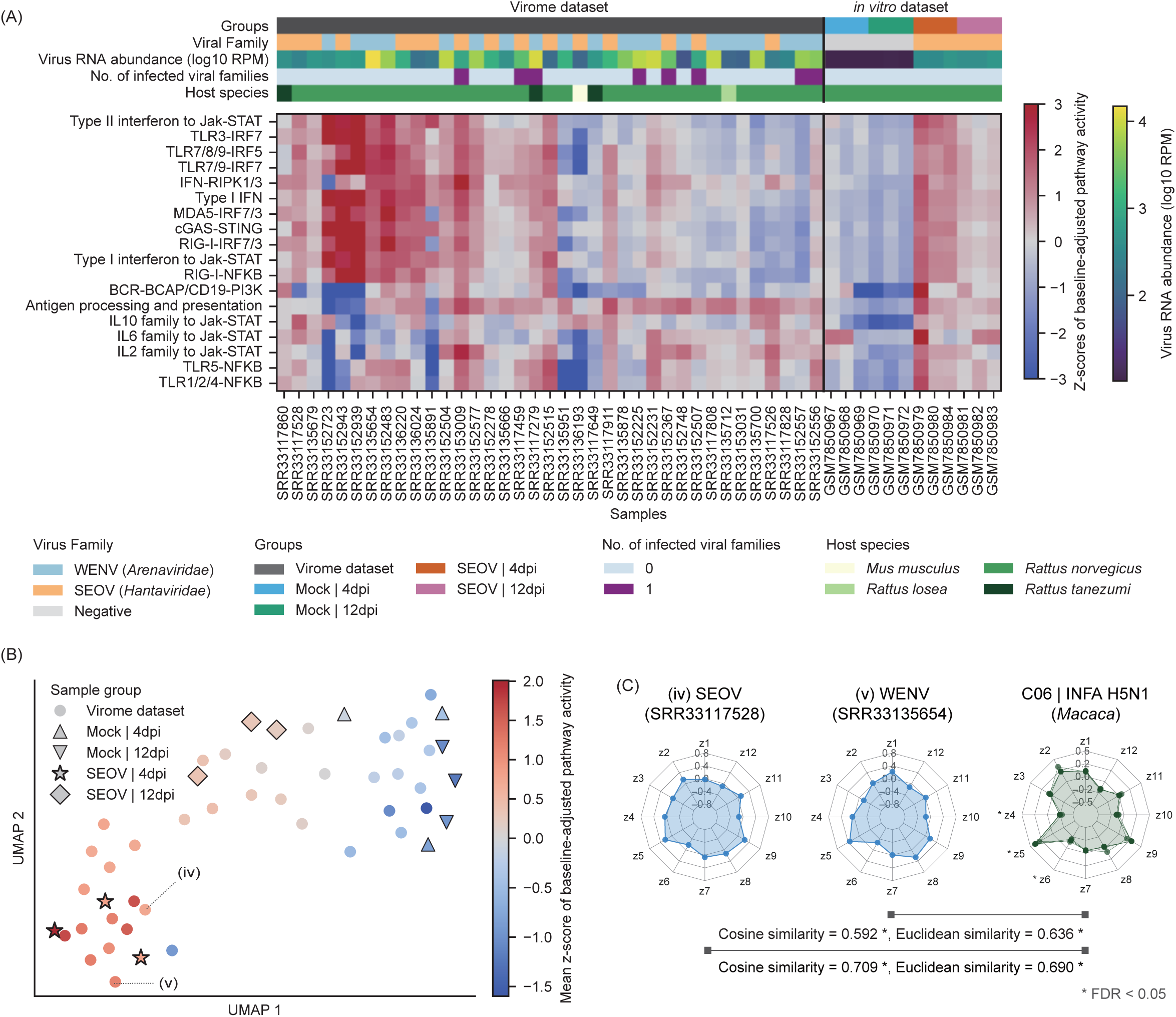
Diversity of host responses to rodent-borne viruses in natural infections. **(A)** Pathway activities related to innate immune and inflammatory responses in rodents infected with SEOV or WENV. The left panel shows samples from the virome data set, and the right panel shows samples from an experimental SEOV infection data set. Activities are shown as z-scores of baseline-adjusted pathway activity. **(B)** UMAP visualization of host response profiles from SEOV- or WENV-infected samples. Points are colored according to the mean z-score of baseline-adjusted pathway activity across the pathways shown in (A). **(C)** Comparisons of host-response profiles between SEOV- or WENV-infected virome samples with high z5 scores and INFA H5N1-infected reference samples assigned to C06. Labels (iv) and (v) correspond to the samples indicated in (B).

## Discussion

We developed ViroChart, a comparative transcriptomic framework for quantitatively profiling host responses to viral infection. By integrating the activities of 484 KEGG pathways into 12 latent factors, ViroChart represented host responses in a common latent space that captured variation associated with the infecting virus and disease progression (**Fig. 1**). We further constructed a reference panel of well-characterized respiratory virus infections and identified shared host-response patterns across taxonomically distinct viruses (**Fig. 2**). Although most samples in virome data sets from wild and farmed animals showed weak host responses (**Fig. 4)**, others exhibited strong inflammatory responses resembling those induced by viruses capable of causing fatal disease (**Figs. 3 and 5**). Together, these results demonstrate that ViroChart provides quantitative host-response information from the same RNA-seq data collected for viral sequence detection, providing additional biological context that may help characterize newly detected viruses and inform their risk assessment. The ViroChart analysis pipeline and pretrained model are publicly available on GitHub (https://github.com/virus-informatics/ViroChart), allowing newly acquired virome RNA-seq data sets to be projected into the existing latent space and compared with host responses from other studies.

Application of ViroChart to animal virome data sets identified particularly strong inflammatory responses associated with two *Betacoronavirus* infections, Bat-CoV HKU5 and RbCoV1 (**Fig. 3C–E**). Although the original studies reported these viral infections, they did not characterize the associated host responses^24^. Notably, these viruses have received markedly different levels of research attention: Bat-CoV HKU5 has been intensively investigated because it can use ACE2 from multiple mammalian species, including humans^24,36–38^, whereas RbCoV1 remains poorly characterized. This contrast illustrates a broader challenge in virome research, in which prior knowledge of potential risk can strongly influence which newly discovered viruses receive further investigation^2,39^. In contrast, ViroChart provides a complementary approach by extracting host-response profiles from all RNA-seq samples in which viruses are detected. Such systematic profiling may help flag viruses of potential concern that would otherwise be overlooked, particularly when prior biological information is limited.

ViroChart also revealed substantial host-response diversity among rodents naturally infected with SEOV or WENV (**Fig. 5**). Although most SEOV-infected animals showed no marked innate immune or inflammatory responses, a subset showed strong responses resembling those observed at early time points after SEOV infection of cell lines. This finding suggests that SEOV, which has been thought to cause inapparent infection in its natural rodent hosts, can elicit such strong responses under certain conditions, possibly early after infection. Interestingly, similarly strong responses were observed in a subset of WENV-infected samples, raising the possibility that WENV infection follows a similar course, with a transient acute phase preceding inapparent infection. Together, these observations illustrate how ViroChart can complement experimental approaches by revealing the distribution of host responses across naturally infected populations—information that controlled infection experiments cannot readily provide.

Although some virus-positive samples in animal virome data sets showed strong inflammatory responses, many showed little or no detectable alteration in their host-response profiles (**Fig. 4**). Several factors could account for these weak responses. Biologically, some viruses may cause low-pathogenicity or asymptomatic infections in their natural hosts, while weak responses may also reflect early stages of infection or viral loads insufficient to elicit substantial host responses. For example, the weak responses observed in influenza virus-positive *Cavia* samples are consistent with previous observations that guinea pigs can support high levels of influenza virus replication without overt symptoms^40,41^. The frequent observation of weak host responses may also reflect differences in the duration of infection stages: the chronic phase generally lasts much longer than the acute phase, potentially resulting in virus-positive samples being dominated by chronically infected animals with weaker responses. Methodological factors may also contribute, as ViroChart estimates infection-associated changes by subtracting the mean response of uninfected samples, which may not adequately account for inter-individual variation in baseline host responses (see below). In addition, sequence contamination, a recognized challenge in virome studies^42^, could result in apparent virus detection without true infection. At present, these possibilities cannot be distinguished, highlighting the need for careful interpretation of virus-positive samples with little evidence of a host response.

This study has several technical limitations. First, the observed host-response profiles may reflect responses to other pathogens, such as bacteria, particularly in virome data sets derived from naturally infected animals. Second, although ViroChart can infer latent factor scores from a single sample, interpreting infection-associated changes requires an appropriate baseline derived from uninfected samples. We used the mean response of uninfected samples from the corresponding host species and tissue as the baseline, which may not capture inter-individual variation and could lead to over- or underestimation of infection-associated changes. Future development should therefore incorporate inter-individual variation into baseline estimation, potentially by leveraging large collections of uninfected RNA-seq data. Third, because ViroChart uses KEGG-defined pathways shared across mammalian species, species-specific biological processes may not be adequately captured. Finally, the latent factors extracted by ViroChart represent statistical covariation among pathway activities and do not necessarily indicate direct biological relationships. In this study, we interpreted each latent factor based on pathways with high loadings, but other biological interpretations may also be possible.

A further consideration is the application of ViroChart to pathogen risk assessment. Although ViroChart captures transcriptomic signatures associated with disease processes, these signatures represent only one component of the broader disease phenotype and do not by themselves indicate viral pathogenicity. In addition, neither strong host responses nor pathogenicity in animals necessarily indicates pandemic potential, as viruses associated with asymptomatic infections in animals have also led to human pandemics^43,44^. Thus, ViroChart should be considered as one component of pathogen risk assessment, complementing other biological and epidemiological evidence.

Despite these limitations, our study demonstrates that host-response profiles can provide complementary biological information from the same RNA-seq data collected for virome analysis. Applying ViroChart to virome RNA-seq data sets could link viral genomic information with quantitative host-response profiles at the level of individual infection samples. Together, such integration could facilitate the systematic characterization and prioritization of newly discovered viruses.

## Methods

### Construction of the ViroChart training data set

We used 5,598 publicly available mammalian RNA-seq samples to train ViroChart, comprising 2,482 virus-infected and 3,116 uninfected samples (**Data set 1**). To capture diverse transcriptional responses associated with viral infection, we used virus detection results from two previous large-scale analyses of public RNA-seq data^7,12^. The first study screened approximately 46,000 RNA-seq samples for viral sequences by performing BLASTx searches against contigs assembled using SPAdes. The second study screened approximately 168,000 RNA-seq samples using geNomad to identify viral sequences in contigs available through the Logan database.

To prioritize samples with well-supported infection status, we primarily selected experimental infection data sets that utilized RNA viruses. For several enteric viruses, including *Sapovirus*, *Kobuvirus*, *Sapelovirus*, *Teschovirus*, *Rotavirus*, and *Torovirus*, we supplemented the training data set with naturally infected samples because experimental infection data sets were limited. Host animal species were selected based on the availability of both gene and pathway annotations in the KEGG database^45–47^, and virus genera with at least five infected samples were included. To reduce overrepresentation of particular host species or research projects, uninfected samples were down-sampled to a maximum of five samples per BioProject and 200 samples per host species. The resulting training data set contained infections with 22 virus genera from 14 virus families across 20 mammalian host species (**Fig. S1A–B**).

### Single-sample pathway analysis using ssGSEA

To summarize gene expression profiles from individual RNA-seq samples at the pathway level, we performed single-sample gene set enrichment analysis (ssGSEA) using the R package GSVA (version 1.50.5)^48,49^. Because ssGSEA calculates gene set scores independently for each sample, gene expression profiles from new RNA-seq samples can be converted into pathway-level activities without comparison to control samples and directly analyzed using the pretrained ViroChart model.

We used KEGG pathways (e.g., map00010) and network modules (e.g., N00001) as gene sets, with the latter representing more specific biological processes. We selected 484 gene sets that were commonly annotated across the mammalian species **(Data set 8)**. Gene sets for each host species were constructed based on KEGG annotations accessed on September 30, 2025 (https://www.genome.jp/kegg/pathway.html).

RNA-seq reads were mapped to the reference genome of each host species using HISAT2 (version 2.2.1)^50^. Reference genomes and gene annotations corresponding to each host species were obtained based on the KEGG organism database (https://www.genome.jp/kegg/tables/br08606.html). Mapped reads were quantified at the gene level using featureCounts (version 2.0.6)^51^ with the corresponding gene annotation, and TPM values were calculated from the resulting gene counts. The TPM expression matrices were then used as input for ssGSEA.

### Bayesian factor analysis

ViroChart uses Bayesian factor analysis to summarize 484 biological pathway activity scores into a lower-dimensional representation of host responses (**Fig. 1A–B**).

For each sample *i*, the standardized pathway activity vector *y_i_* was represented by a *K*-dimensional latent factor vector *z_i_*. Here, *W* denotes the loading matrix relating latent factors to pathway activities, and ψ represents pathway-specific observation noise. The model was formulated as follows:

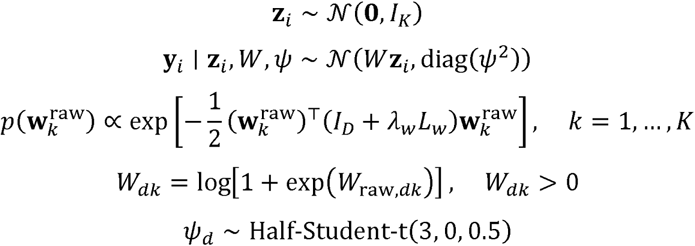

To facilitate interpretation of latent factor direction, we constrained the loading matrix *W* to be non-negative. Under this constraint, higher scores for a given latent factor correspond to higher activities of pathways with large loadings on that factor. Specifically, we estimated an unconstrained loading parameter *W*_raw_ and applied a softplus transformation to obtain the non-negative loading matrix *W*.

Because pathways with similar gene composition are likely to show coordinated activity patterns, we further introduced graph Laplacian regularization on *W*_raw_. This regularization encouraged pathways with similar gene composition to have similar loadings on the same latent factors. The pathway graph was constructed based on Jaccard similarities between gene sets corresponding to human KEGG pathways and network modules.

Before model fitting, the ssGSEA score for each pathway was standardized to have a mean of 0 and a standard deviation of 1 across the training data set. When projecting new samples into the pretrained model, the same centering and scaling parameters estimated from the training data set were applied. The model was fitted using CmdStanR (version 0.8.0; https://mc-stan.org/cmdstanr/) with CmdStan (version 2.38.0; https://github.com/stan-dev/cmdstan). Maximum a posteriori (MAP) estimates were obtained by numerical optimization using the L-BFGS algorithm with a maximum of 2,000 iterations. After model fitting, the latent factor scores *z* were extracted for each sample and used as features in a *k*NN classifier (*k* = 10) to predict the infecting virus genus. The dimensionality of the latent space was evaluated across different values of *K* based on *k*NN classification performance using the resulting latent factor scores and five-fold cross-validation, as described below (**Fig. S2E**). Because classification performance largely plateaued at *K*=12, we selected *K*=12 for the final ViroChart model (**Fig. S1C**).

### Evaluation of model projection performance

To evaluate the ability of ViroChart to project new RNA-seq samples into a pretrained latent space, we constructed five hold-out models using five-fold stratified group cross-validation, with BioProject used as the grouping unit (**Fig. S2**). Samples were partitioned while balancing the distribution of virus genera across folds as much as possible and ensuring that samples from the same BioProject were not included in both the training and evaluation sets. Cross-validation was implemented using scikit-learn (version 1.7.2)^52^.

For each hold-out model, all trained model parameters were fixed, and only the latent factor scores z of the evaluation samples were estimated by MAP estimation. Projection performance was evaluated from four perspectives. First, to assess model fit, we calculated the log likelihood of each evaluation sample and compared its distribution with that of the training samples in the full model. Second, to assess preservation of the global structure of the latent space, we calculated pairwise Euclidean distances between samples and assessed their Spearman correlation with the corresponding distances obtained from the full model. Third, preservation of local neighborhood structure was evaluated by identifying the *k* nearest neighbors of each sample (*k* = 10, 50, and 100) and calculating the Jaccard similarity between the neighbor sets obtained from each hold-out model and the full model. Finally, *k*NN classifiers were constructed using the latent factor scores as features, and classification performance was evaluated for infection status and infecting virus genus on evaluation samples. The number of neighbors was evaluated at *k* = 1, 3, 5, 10, and 20, and the value yielding the best classification performance was selected. The value yielding the highest mean macro-F1 score across cross-validation folds was selected, resulting in *k* = 20 for infection-status classification and *k* = 1 for virus-genus classification.

To assess how differences in the training data set affected the interpretability of latent factors, we compared the loading matrices W between the full model and each hold-out model. Because the ordering of latent factors is not inherently identifiable across independently fitted models, we calculated Spearman correlation coefficients between all pairs of loading vectors from the full and hold-out models to generate a K × K correlation matrix. We then used the Hungarian algorithm to identify the one-to-one factor matching that maximized the sum of the absolute correlation coefficients. For each matched factor pair, similarity was evaluated using the Spearman correlation between loading vectors and the Jaccard index of the top 50 pathways ranked by loading.

### Analysis of ViroChart-derived host-response profiles

Latent factor scores were obtained differently depending on the source of the samples. For samples in the training data set, latent factor scores obtained during model fitting were used (**Figs. 1–2**). For samples derived from virome data sets, latent factor scores were estimated by projecting each sample into the pretrained model while fixing all model parameters (**Figs. 3–5**).

UMAP was performed using the Python package umap-learn (version 0.5.9)^53^ to visualize variation in ViroChart-derived host-response profiles. The UMAP shown in **Fig. 1C–D** was constructed directly from the estimated latent factor scores. Because these scores were substantially influenced by host species and tissue, we defined a baseline-adjusted host-response profile for each sample to better represent infection-associated changes. Specifically, the mean latent factor scores of uninfected samples from the corresponding host species and tissue were subtracted from the latent factor scores of each sample. The UMAP in **Fig. 2A** was constructed from these baseline-adjusted host-response profiles, and virome samples were projected onto the same UMAP space (**Fig. 3A–B**). All radar plots were also generated using the baseline-adjusted host-response profiles.

PERMANOVA was used to evaluate the contribution of sample metadata to variation in latent factor scores (**Fig. 1C–D**) or baseline-adjusted host-response profiles (**Fig. 3A**). Euclidean distance matrices were calculated from the corresponding scores or profiles and analyzed using the adonis2 function in the R package vegan (version 2.7-3; https://cran.r-project.org/web/packages/vegan/index.html). In **Fig. 1C**, the contribution of infecting virus genus was evaluated after accounting for host genus and tissue; in **Fig. 1D**, the contribution of days post-infection (dpi) was evaluated after accounting for BioProject and infection condition (the route and dose of virus inoculation). For the analysis in **Fig. 3A**, we restricted samples to those infected with SEOV or WENV and evaluated the contribution of each of the following variables to host response profiles: infecting virus species, viral RNA abundance, the number of coinfecting virus families, and host species. When covariates were included, they were entered into the model before the variable of interest using sequential tests. Statistical significance was assessed using 999 permutations.

### Host-response similarity analysis

Host-response similarity was evaluated using two complementary metrics based on baseline-adjusted host-response profiles (**Figs. 2C–E, 3C–E, and 5C**). Cosine similarity quantified similarity in the direction of host responses, whereas Euclidean-distance-based similarity quantified their overall proximity in the 12-dimensional latent space. Euclidean-distance-based similarity was defined as 1/(1+d), where d represents the Euclidean distance between two profiles.

To assess whether the observed similarity was greater than expected by chance, we performed query-fixed permutation tests with 10,000 permutations. For cluster-to-cluster comparisons, target samples were randomly selected from the full sample set while preserving the size of the observed target cluster. The mean similarity between the query and randomized target clusters was calculated to generate an empirical null distribution. For sample-to-cluster comparisons, sets of samples matching the size of the target cluster were randomly selected from all samples except the query sample. The mean similarity between the query sample and each randomized target set was used to generate the null distribution. For sample-to-sample comparisons, a single target sample was randomly selected from all samples except the query sample in each permutation. Empirical *P* values were calculated from the corresponding null distributions and adjusted for multiple testing using the Benjamini–Hochberg procedure.

### Analysis of host responses to respiratory virus infections

To compare host responses to respiratory virus infections, we collected 317 lung RNA-seq samples from five animal species infected with four respiratory viruses (**Data set 4**). The abundance of viral RNA in each sample was estimated from publicly available NCBI SRA Taxonomy Analysis Tool (STAT) results retrieved through the NCBI SRA BigQuery tables^54^. For each sample, we extracted the number of reads assigned to the infecting virus and normalized this value by the total number of reads.

To examine the diversity of host-response profiles, k-means clustering was performed using the baseline-adjusted host-response profiles (**Fig. 2A**). The number of clusters was evaluated from *k*=8 to 18 based on the mean silhouette coefficient, resulting in *k*=16.

To compare changes in innate immune and inflammatory pathway activities across samples, pathway activity scores were baseline-adjusted by subtracting the mean scores of uninfected samples from the corresponding host genus and tissue category. The adjusted scores were then converted to robust z-scores separately for each pathway using all available samples from the same host genus as the reference distribution, irrespective of infection status (**Fig. 2F**).

### Analysis of host responses in virome data sets

We analyzed 1,002 samples from four BioProjects as virome data sets. For the analyses in **Fig. 3**, we used 199 lung samples from two BioProjects (PRJCA026706 and PRJNA1238520)^24,25^. For **Fig. 4**, we expanded the analysis to all tissues and included two additional bat BioProjects (PRJNA1184956 and PRJNA929070)^27,28^. We used publicly available virus detection results from previous virome studies that analyzed these data sets. Latent factor scores for each sample were estimated by projection into the pretrained ViroChart model.

For **Fig. 3A–B**, virome samples were projected onto the UMAP space constructed in **Fig. 2A** to compare their host-response profiles with those of well-characterized respiratory viruses. For **Figs. 3B, 5A, and S4A,** pathway activity scores or latent factor scores were baseline-adjusted and converted to robust z-scores as in **Fig. 2F**. For the SEOV experimental samples in **Fig. 5A**, baseline adjustment used mock samples, whereas robust z-score standardization used all mock and infected samples from the same BioProject. For **Fig. 5B**, we constructed a new UMAP using the samples and pathways shown in **Fig. 5A**, with each point colored by its mean pathway activity score.

For **Fig. 3F–G**, we compared pathway activities between clusters C06 and C07 and between two Bat-CoV HKU5-positive samples assigned to C06-like and C07-like profiles. The C06–C07 difference was estimated using a limma linear model with empirical Bayes moderation^55^, whereas the difference between the HKU5-positive samples was calculated as the pathway activity in the C06-like sample minus that in the C07-like sample. We ranked pathways by the absolute value of their z5 loadings and used a one-sided Fisher’s exact test to assess whether the top 20 were enriched among pathways with a positive C06–C07 difference (Benjamini–Hochberg FDR < 0.05) and a positive difference between the two HKU5-positive samples.

For SEOV- or WENV-infected samples, we used Centrifuger^56^ to assess read composition and verify consistency with the registered host information. The results indicated that nearly all samples originated from *Rattus*, whereas one sample was assigned to *Mus musculus* (**Data set 7**).

### Host-response intensity analysis in the virome data set

For **Fig. 4A and 4C**, host-response intensity was defined as the sum of the absolute values of the 12 dimensions of the baseline-adjusted host-response profile 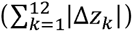 Host-response intensity was then converted to a robust z-score using uninfected samples from the same host genus as the reference distribution. We refer to these scores as uninfected-reference z-scores of response intensity.

For **Fig. 4A**, we fitted a linear regression model including viral RNA abundance, host taxonomic group, and infecting virus family as explanatory variables to evaluate the overall association between viral RNA abundance and host-response intensity. To determine whether this association differed among host taxonomic groups, we fitted an interaction model that additionally included an interaction between viral RNA abundance and host taxonomic group. For these regression models, we included samples with viral RNA abundance >10 RPM and used log10(RPM + 1) as the explanatory variable. The overall interaction was evaluated using a Wald test. Group-specific regression coefficients and two-sided P values were estimated from the interaction model.

For **Fig. 4C**, to compare response intensity across virus–host combinations, each combination was defined by the infecting virus family and host genus. We analyzed virus-positive samples with viral RNA abundance >10 RPM and included combinations represented by at least five samples. We fitted a linear regression model including virus–host combination and viral RNA abundance centered at its median as explanatory variables. Heteroscedasticity-consistent standard errors (HC3) were used. Significantly positive adjusted response intensities were identified using one-sided tests against zero, with Benjamini–Hochberg correction (FDR < 0.05).

In **Fig. 4B** and **Fig. S4B**, we examined host responses at the pathway level. As in **Fig. 4A**, pathway activity was expressed as an uninfected-reference z-score for each pathway. For **Fig. 4B**, we applied the interaction model described for **Fig. 4A** to each pathway separately. We estimated host-group-specific slopes using HC3 standard errors and adjusted their two-sided P values across all pathway–host-group comparisons using the Benjamini–Hochberg method.

For **Fig. S4B**, we grouped the selected pathways into “Nucleic acid sensing–IRF/IFN signaling” and “Cytokine-mediated immune regulation.” We fitted a mixed-effects model with an interaction among viral RNA abundance, host taxonomic group, and pathway category, adjusting for infecting virus family. The model included a random intercept for each sample and random intercepts and slopes for each pathway. Category-level regression lines, 95% confidence intervals, slopes, and two-sided P values were derived from the fixed effects.

## Data Availability

The data sets and scripts used in this study are available through the Zenodo repository (https://doi.org/10.5281/zenodo.23201228)^57^.

## Code Availability

The source code used in this study and the ViroChart package are publicly available on GitHub (https://github.com/virus-informatics/ViroChart).

## Data sets

Data set 1: ViroChart training data set.

Data set 2: Loading vectors for the ViroChart latent factors.

Data set 3: Experimental Ebola virus infection data set.

Data set 4: Respiratory virus infection data set.

Data set 5: Animal virome data set used for lung-specific analyses.

Data set 6: Animal virome data set including samples from multiple tissues.

Data set 7: Virome data set of wild rodents infected with hantavirus and arenavirus.

Data set 8: Gene sets used for pathway activity analysis.

## Acknowledgments

We thank Professor Shuo Su, Yuqi Shi, and their colleagues for their generous assistance with data sharing, although the data were not used in this study. We also thank Dr. Arnon Plianchaisuk and Dr. Amjad Khalaf for their constructive feedback on the manuscript development. Supercomputing resources were provided by the Human Genome Center, The Institute of Medical Science, The University of Tokyo. We thank YouWorks Inc. and Holonics Inc. for their technical support in the development and implementation of the ViroChart software package. BioRender was used to create selected graphical elements in the figures.

This study was supported in part by JST PRESTO (JPMJPR23R4, to J.K.); JSPS KAKENHI (JP25H01310, JP25H01309, JP26K18192 to J.K.); cSIMVa Vaccine Challenge Grant (to J.K.); the Inohana Foundation (Chiba University) Grant-in-Aid (IFCU-2025-03) (to J.K.); JST, Moonshot R&D (JPMJMS2025, to T.S.); the Japan Agency for Medical Research and Development (AMED) (JP223fa627003, JP263f0134003, JP25fk0108732, JP25fk0108656 to T.S.); the Emerging and Re-emerging Infectious Diseases and Vaccination Policy Promotion Research Project of the Ministry of Health, Labour and Welfare, Japan (26HA2015 to T.S.); AMED SCARDA Center for Advanced Modalities and Drug Delivery Systems “CAMaD” (JP223fa627002 and JP253m0134002, to J.I.); AMED Japan Program for Infectious Diseases Research and Infrastructure (Interdisciplinary Cutting-edge Research) (JP26wm0325087, to J.I.); AMED-CREST (JP26gm2010010, to J.I.); and JSPS KAKENHI Grant-in-Aid for Scientific Research B (JP25K00116, to J.I.).

## Author Contributions

Project design, management and supervision: Junna Kawasaki, Jumpei Ito, and Tadaki Suzuki

Development of the ViroChart framework: Junna Kawasaki

Collection and curation of data sets: Junna Kawasaki

Bioinformatics analyses: Junna Kawasaki

ViroChart package development: Junna Kawasaki

Data visualization and figure preparation: Junna Kawasaki

Support for bioinformatics analyses: Michiaki Hamada and Luca Nishimura

Manuscript drafting: Junna Kawasaki

Manuscript editing: Junna Kawasaki, Jumpei Ito, and Edward C. Holmes

All authors reviewed and approved the final manuscript.

---

**Fig. S1.**
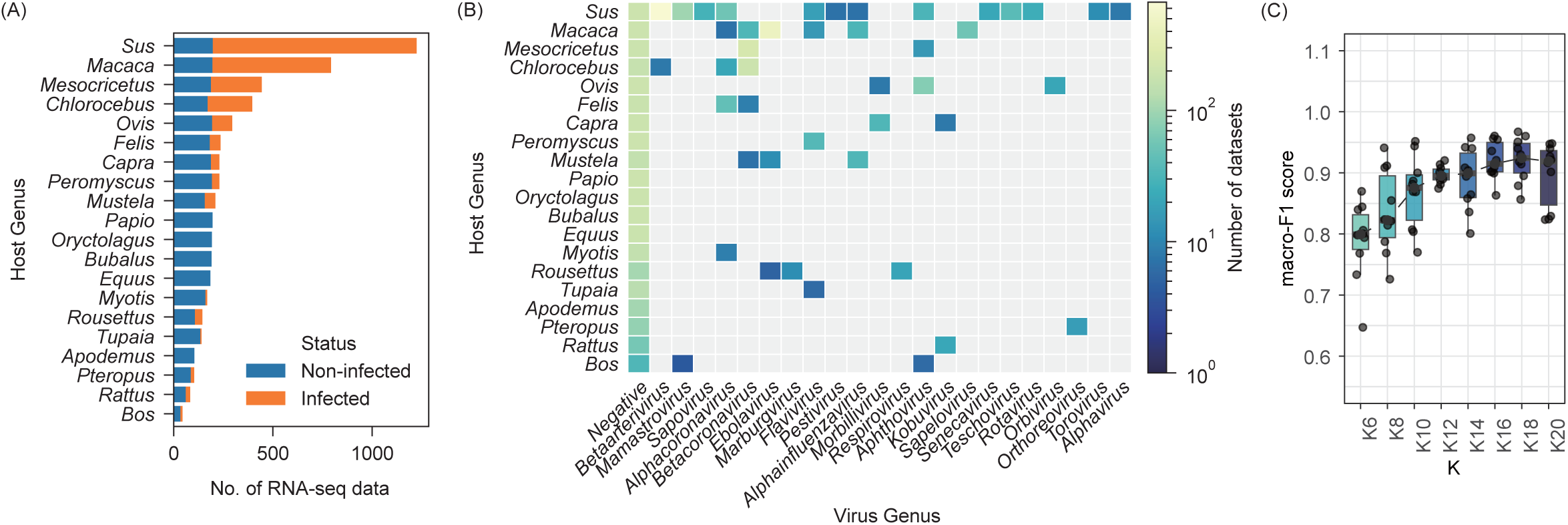
Training of ViroChart. **(A)** Proportions of virus-infected and uninfected samples for each host species in the ViroChart training data set. **(B)** Numbers of samples according to infecting virus genus and host animal genus in the ViroChart training data set. **(C)** Performance of *k*-nearest neighbors (*k*NN; *k* = 10 neighbors) classification of the infecting virus genus across different numbers of latent factors, evaluated using five-fold cross-validation.

**Fig. S2.**
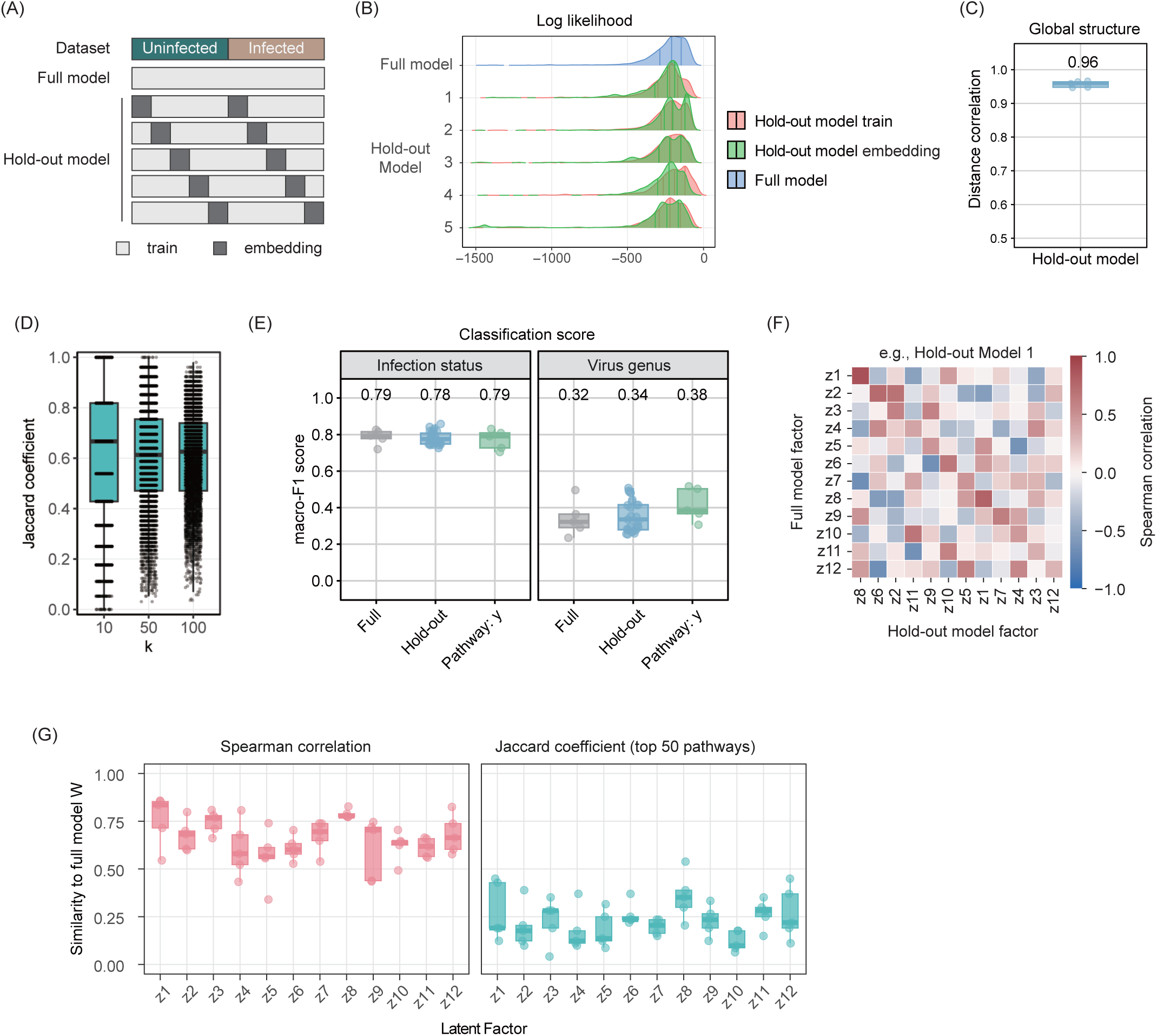
Evaluation of ViroChart projection capability and robustness of latent factor interpretation. **(A)** Overview of the hold-out models. Five hold-out models were generated by excluding subsets of BioProjects from the training data set. Samples excluded from each training data set were projected into the corresponding latent space. **(B)** Log likelihood values for training samples in the full model and projected samples in the hold-out models. **(C)** Correlation of pairwise Euclidean distances between samples in the latent spaces inferred by the full and hold-out models. **(D)** Similarity of the *k*-nearest neighbor structure of projected samples between the full and hold-out models (*k* = 10, 50, and 100). **(E)** *k*NN classification performance for infection status (*k* = 20) and infecting virus genus (*k* = 1) using latent factor scores from the full and hold-out models. Performance was evaluated across the five cross-validation folds. The Pathway (y) model uses 484 pathway activity scores as features. **(F)** Correlations between latent factor loading vectors in the full and hold-out models. One of the five hold-out models is shown as an example. **(G)** Similarity of latent factor loading vectors between the full and hold-out models after factor matching.

**Fig S3.**
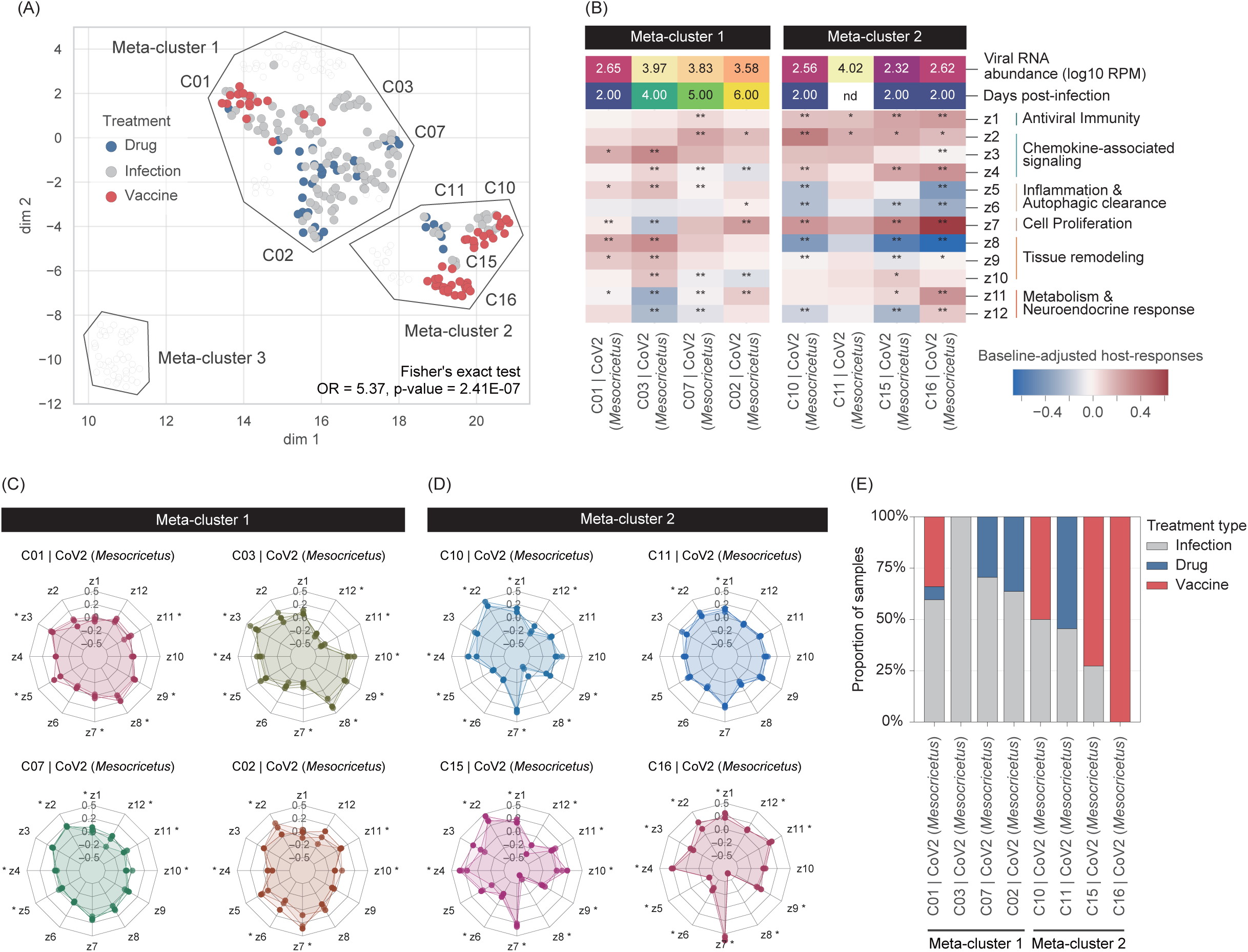
Characteristics of host response clusters associated with SARS-CoV-2 infected hamsters. **(A)** UMAP visualization based on baseline-adjusted host-response profiles of respiratory virus-infected samples. Points are colored according to treatment type. Enrichment of therapeutic drug-treated or vaccinated samples in Meta-cluster 2 was assessed using Fisher’s exact test. **(B)** Mean baseline-adjusted latent factor scores for clusters containing SARS-CoV-2-infected hamster samples. Asterisks indicate latent factors that differed significantly from those in uninfected samples (Welch’s t-test with Benjamini–Hochberg correction; *FDR < 0.05, **FDR < 0.01). Viral RNA abundance and days post-infection (dpi) are shown as medians for each cluster. **(C-D)** Representative baseline-adjusted host-response profiles from Meta-cluster 1 (C) and Meta-cluster 2 (D). **(E)** Proportions of drug-treated or vaccinated samples in each cluster.

**Fig S4.**
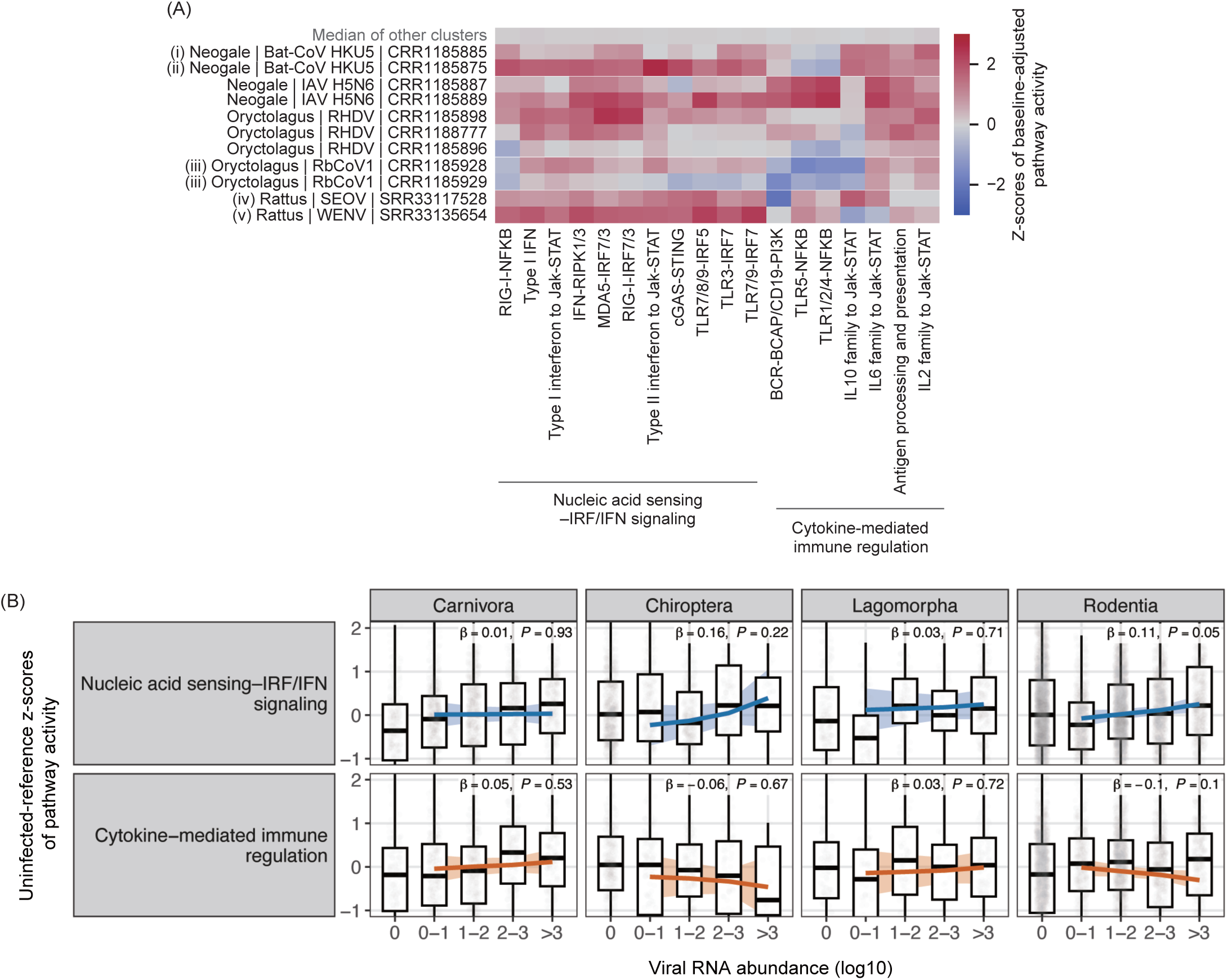
Pathway-level host responses in animal virome data sets. **(A)** Heatmap of pathway activities related to innate immune and inflammatory responses. Values are shown as z-scores of baseline-adjusted pathway activity, using the same calculation method and pathway set as in Fig. 2F. Labels (i)–(v) correspond to the samples indicated in Figs. 3 and 5. **(B)** Associations between viral RNA abundance and pathway activities, grouped into functional categories. Pathway-specific uninfected-reference z-scores were analyzed using a mixed-effects regression model. Colored lines represent the estimated regression slopes for each functional category, with shaded areas indicating 95% confidence intervals. β and P indicate the estimated category-level slope and its two-sided P value, respectively.

